# Hypothesis-free identification of modulators of genetic risk factors

**DOI:** 10.1101/033217

**Authors:** Daria V. Zhernakova, Patrick Deelen, Martijn Vermaat, Maarten van Iterson, Michiel van Galen, Wibowo Arindrarto, Peter van ‘t Hof, Hailiang Mei, Freerk van Dijk, Harm-Jan Westra, Marc Jan Bonder, Jeroen van Rooij, Marijn Verkerk, P. Mila Jhamai, Matthijs Moed, Szymon M. Kielbasa, Jan Bot, Irene Nooren, René Pool, Jenny van Dongen, Jouke J. Hottenga, Coen D.A. Stehouwer, Carla J.H. van der Kallen, Casper G. Schalkwijk, Alexandra Zhernakova, Yang Li, Ettje F. Tigchelaar, Marian Beekman, Joris Deelen, Diana van Heemst, Leonard H. van den Berg, Albert Hofman, André G. Uitterlinden, Marleen M.J. van Greevenbroek, Jan H. Veldink, Dorret I. Boomsma, Cornelia M. van Duijn, Cisca Wijmenga, P. Eline Slagboom, Morris A. Swertz, Aaron Isaacs, Joyce B.J. van Meurs, Rick Jansen, Bastiaan T. Heijmans, Peter A.C. ‘t Hoen, Lude Franke

## Abstract

Genetic risk factors often localize in non-coding regions of the genome with unknown effects on disease etiology. Expression quantitative trait loci (eQTLs) help to explain the regulatory mechanisms underlying the association of genetic risk factors with disease. More mechanistic insights can be derived from knowledge of the context, such as cell type or the activity of signaling pathways, influencing the nature and strength of eQTLs. Here, we generated peripheral blood RNA-seq data from 2,116 unrelated Dutch individuals and systematically identified these context-dependent eQTLs using a hypothesis-free strategy that does not require prior knowledge on the identity of the modifiers. Out of the 23,060 significant *cis*-regulated genes (false discovery rate ≤ 0.05), 2,743 genes (12%) show context-dependent eQTL effects. The majority of those were influenced by cell type composition, revealing eQTLs that are particularly strong in cell types such as CD4+ T-cells, erythrocytes, and even lowly abundant eosinophils. A set of 145 *cis*-eQTLs were influenced by the activity of the type I interferon signaling pathway and we identified several *cis*-eQTLs that are modulated by specific transcription factors that bind to the eQTL SNPs. This demonstrates that large-scale eQTL studies in unchallenged individuals can complement perturbation experiments to gain better insight in regulatory networks and their stimuli.

## Introduction

The molecular mechanisms underlying the association of genetic risk factors with disease and complex traits are still largely elusive. Many disease-associated genetic variants are found in non-coding parts of the genome ^1,2^ and thus must have a regulatory effect on expression. Mapping single nucleotide polymorphisms (SNPs) with an effect on the regulation of gene expression (expression quantitative trait loci, eQTLs) helps to unravel the regulatory networks that underlie physiological traits and diseases ^3–8^. Given differences between the regulatory networks of different cell types, it is not surprising that a substantial fraction of eQTLs are only apparent in specific cell types or tissues ^9–14^. The presence of external stimuli and the activity of internal signaling pathways may also determine the presence and strength of the regulatory effects of eQTLs. For example, a subset of eQTLs in immune cells may only be observed after activation of these cells by immunological triggers ^15–20^. Knowledge of the cellular context in which disease-associated eQTLs are active can help to identify the cell types that are relevant in the pathophysiology; identification of the cell type in which a risk locus shows the most profound effects allows prioritization of variants for functional experiments. Additionally, insights into the activity of signaling pathways modifying eQTL effects help to unravel the regulatory networks underlying disease. Here, we developed and applied a strategy to identify the most important intrinsic and extrinsic factors that modify eQTL effects in blood cells, without making any prior assumptions on the identity of these modifiers. We demonstrate how the eQTLs and their modifiers contribute to better understand the molecular basis of disease.

## Results

### Main-effect *cis*-eQTLs

We generated a comprehensive set of *cis*-eQTLs by sequencing whole peripheral blood mRNA of 2,176 healthy adults from four Dutch cohorts ^21–24^ (2,116 individuals remaining after stringent quality control (Table S1, Supplementary material)). We quantified gene and exon expression, as well as exon ratios (the proportion of expression of an exon relative to the total expression of all exons of a gene) and polyA ratios (the ratio of the expression in upstream and downstream parts of the 3’-UTRs separated by annotated polyadenylation (polyA) sites) and performed *cis-* eQTL mapping for all of these. We detected *cis*-eQTL effects for 66% of the protein coding genes tested and 19% of the non-coding genes tested. In total, we found eQTL effects for 23,060 different genes (false discovery rate (FDR) ≤ 0.05). We replicated 84% of 6,418 previously reported *cis*-eQTL genes that we had previously detected in a meta-analysis of 5,311 array-based blood samples ^4^ (90% with the same allelic direction) (Table S2). This demonstrates the superior statistical power to detect eQTLs when using RNA-seq data (Tables S2 and S3). We also observed strong overlap with RNA-seq based *cis*-eQTLs from EBV-transformed lymphoblastoid cell lines (LCL) ^5^ (78% of the LCL *cis*-eQTLs could be replicated, 88% with the same allelic direction), but substantially extended the list of genes that are known to be under genetic regulation (replication results in Supplementary material online, Table S2). In addition to detected gene-level eQTLs, we identified for 21,888 different genes with one or more exon-level QTL effects and 9,777 and 2,322 genes where SNPs affected the inclusion rate of exons and the usage of polyA sites, respectively (Table S3). A complete catalogue of all our eQTLs can be downloaded and explored via a dedicated browser at http://genenetwork.nl/biosqtlbrowser.

Multiple unlinked SNPs in the same locus may independently influence expression or mRNA processing of the same gene ^25^. We analyzed this using stepwise regression of the effects of the top eQTL SNPs. More than half of the *cis*-regulated genes showed evidence for multiple independent eQTL effects (Figure 1a, Figure S1).

**Figure 1.**
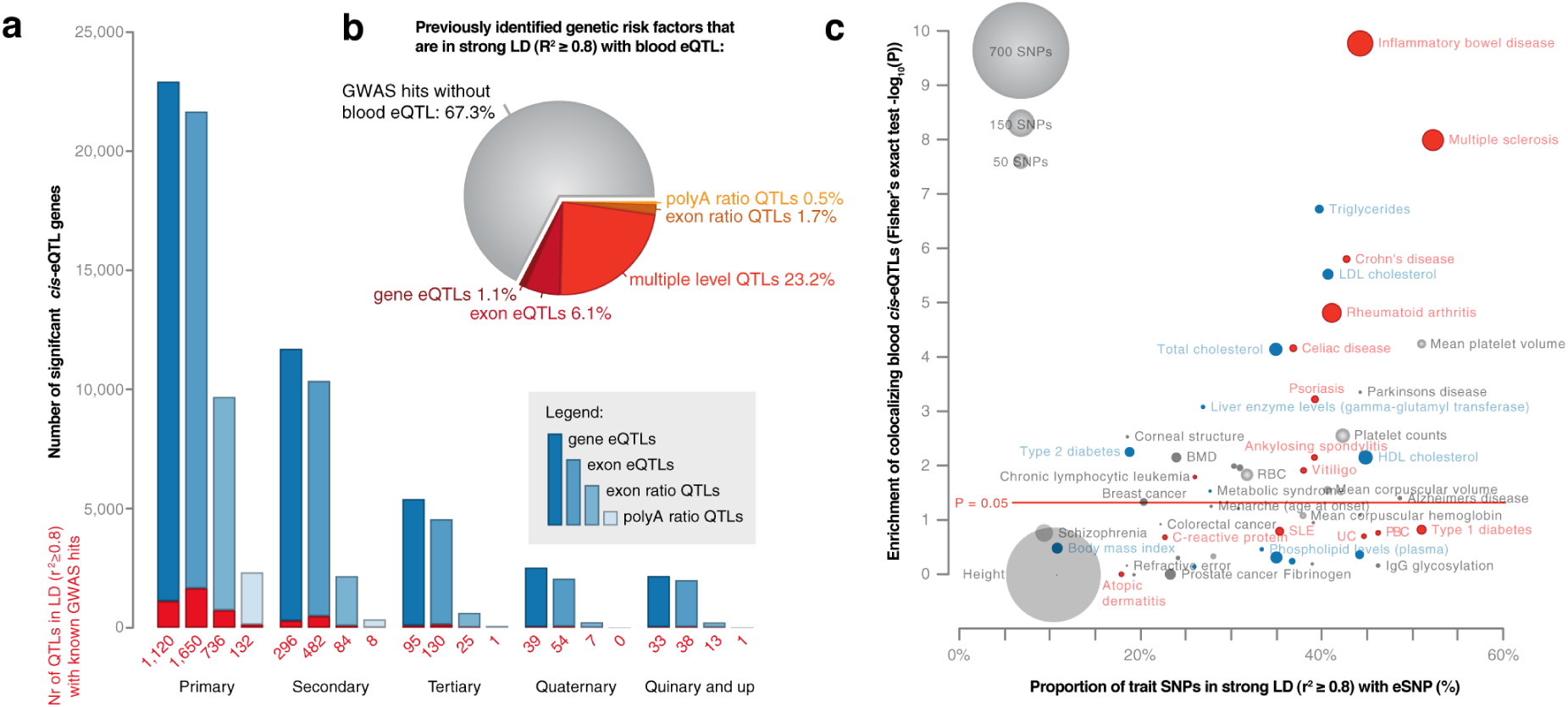
Over 20,000 genes are regulated by *cis*-eQTLs overlapping with 33% of the entries in the GWAS catalog. (A) Bar plot listing the numbers of *cis*-regulated genes having one, two, three, four and five or more independent eQTL effects (FDR ≤ 0.05). Different shades of blue represent the type of eQTL (gene, exon, exon ratio or polyA ratio). The number of eQTLs overlapping with SNPs in the GWAS catalog (r^2^ ≥ 0.8) are indicated in red. (B) Distribution of GWAS catalog variants over the different types of eQTLs. Of the GWAS catalog SNPs, 8% affect exon-level QTLs or polyA ratio QTLs but not overall gene expression levels. (C) Auto-immune disorders and traits related to blood show a higher co-localization with eQTLs compared to anthropometric traits and diseases without an immune or hematological component.

The gene *cis*-eQTL SNPs are strongly enriched for DNase I footprints, various histone marks and binding sites of multiple transcription factors ^26^ (Table S4) suggesting that our substantial sample-size enabled us to pinpoint likely causal regulatory variants. Moreover, top eQTL SNPs were significantly enriched for general and blood-cell-type-specific enhancers (as taken from Andersson et al., 2014 ^27^), but not for non-blood tissue-specific enhancers (Table S5). Evidence for the functionality of exon ratio and polyA ratio QTLs in mRNA splicing and polyadenylation is presented in the supplementary material.

One third (2,064 / 32.7%) of previously established genetic risk factors for disease or complex traits (derived from the NHGRI GWAS catalog and a set of reported ImmunoChip associations, P ≤ 5 × 10^−8^) were in strong linkage disequilibrium (LD r^2^ ≥ 0.8) with a top eQTL SNP (Table S6, Figure 1b). As expected, eQTL effects were predominantly found for SNPs associated with hematological, lipid or immune-related traits. We observed a highly significant enrichment of co-localization of eQTL and GWAS SNPs (LD r^2^ ≥ 0.8) for many immune disorders, as compared to height (see supplementary material for details), indicating that our blood *cis*-eQTLs are highly informative for diseases such as inflammatory bowel disease, multiple sclerosis and rheumatoid arthritis (Figure 1c).

### Context-dependent eQTLs

The effects of SNPs on gene expression often depend on the cell type or tissue under investigation ^9–12^, and may be modified by external and environmental factors ^15–19^. To identify these context-dependent eQTLs, we developed a hypothesis-free strategy that does not assume any prior knowledge about the factors that may modify the eQTL effects (Figure 2a). Instead of using known factors, such as the percentage of neutrophils in blood ^28^, in a gene by environment interaction model, we used the expression levels of other genes as interaction factors. Here, we expect that the genes whose expression levels modify eQTLs are proxies of cell types or other intrinsic or extrinsic factors, and we call these genes ‘proxy genes’.

**Figure 2.**
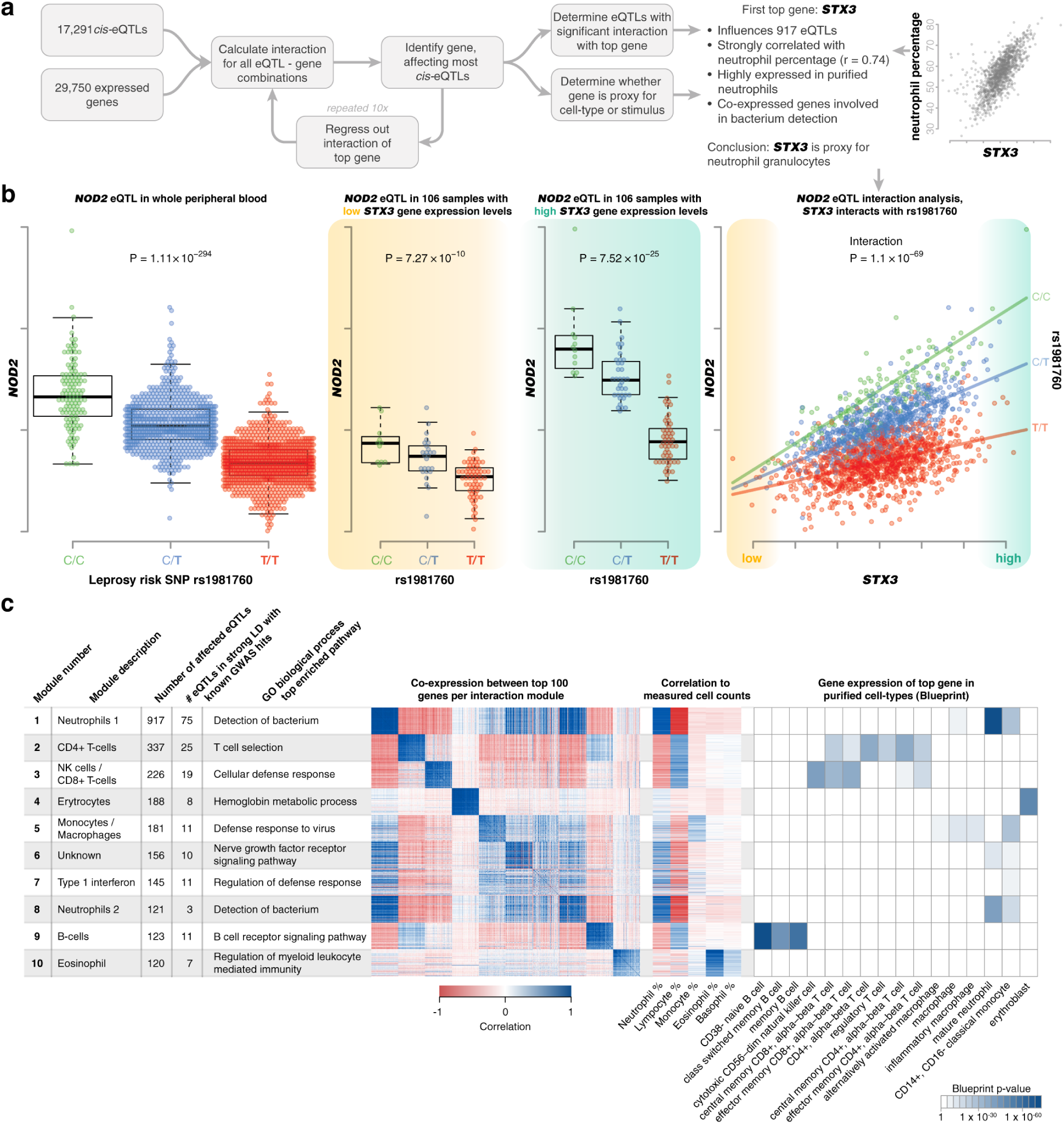
Identification of the strongest modifiers of eQTL effects. (A) Strategy for hypothesis-free identification of modifiers of eQTL effects: each of the highly expressed genes (at least one read in all samples) was tested for its ability to modify each of the 17,291 eQTL effects (for eQTL genes with at least one read in all samples). For each of these covariate genes, we determined the overall strength of the interaction effects with all eQTLs. We selected the strongest covariate gene and regressed-out the interaction term from the data. We did this for 10 iterations allowing the identification of 10 independent modules that affect the strength of eQTLs. (B) An example of a context-dependent eQTL effect is rs1981760, a strong eQTL for the *NOD2* gene. This SNP is in strong LD (r^2^ = 0.99) with rs9302752, a variant associated to Leprosy susceptibility. The leprosy risk allele results in decreased expression of *NOD2*. In samples with low *STX3A* expression, only a weak eQTL effect is observed. In samples with high *STX3A* expression, a strong eQTL effect is observed. In accordance with this, using this *STX3A* gene as a covariate in an interaction model reveals a very strong interaction effect. This *STX3A* gene is the top covariate of our first interaction module. This module correlates strongly to neutrophil percentage (Pearson r = 0.72) and gene enrichment analysis shows involvement in antibacterial response. Furthermore, individuals carrying the Leprosy risk allele have significantly weaker *NOD2* up-regulation in neutrophils compared to non-carriers. This is in line with earlier reports showing this eQTL to be stronger in FACS-sorted neutrophils compared to monocytes ^47^. (C) We annotated each of our 10 modules using the top 100 proxy genes and show that these top 100 genes are strongly correlating per module. This top 100 was used for gene function enrichments (for full results see Table S9) and was correlated to known cell proportions. We used BLUEPRINT expression data for sorted populations of blood cells to validate cell-type-specific expression in each module.

To do this in a systematic manner, we ran interaction analyses for all the eQTLs identified for each gene, evaluating each gene for its potential to influence eQTLs and quantifying per gene the extent to which it influenced the complete set of *cis*-eQTLs. We first identified the proxy gene acting on the highest number of eQTLs. We subsequently corrected the expression data for interaction effects with the top proxy gene and repeated this process in an iterative fashion to find additional independent proxy genes (Figure 2a). We first concentrated on the top 10 proxy genes that independently affected, in total, 1,842 unique *cis*-eQTL genes (Figure 2c).

An example is shown in Figure 2b. We found a significant eQTL effect of SNP rs1981760 (a SNP associated with leprosy susceptibility) on *NOD2* expression. The first top proxy gene, *STX3*, had a significant interaction effect on this eQTL. Samples with very low expression of *STX3* showed only a very weak eQTL on *NOD2*, whereas samples with very high *STX3* expression showed a stronger eQTL effect size. When visualizing this using an interaction plot, a clear divergence in *NOD2* expression between samples with different genotypes can be observed (Figure 2b). Further analysis demonstrated that *STX3* expression was strongly correlated (Pearson r = 0.74) with the percentage of neutrophils in the blood, suggesting that *STX3* is a proxy for neutrophils. In this example, the eQTL effect is found to be more prominent in neutrophils than in other blood cell types, and the expression of *NOD2* found to be lower in carriers of the risk allele compared to carriers of the protective allele. It also confirms a role for neutrophils in the disease for which a genetic association with the locus is found, *i.e*. leprosy.

To investigate the biological context reflected by the 10 independent proxy genes, we selected, for each of these genes, the top 100 genes showing a nearly identical interaction effect with eQTLs (see Methods and Table S7 for full details). As expected, these genes are highly co-expressed (Figure 2c), and we refer to such a set of genes as an ‘interaction module’.

We first assessed whether these interaction modules might represent markers for specific cell types, and correlated the top 100 genes per module with the blood cell counts measured in our samples (neutrophils, lymphocytes, eosinophils, basophils and monocytes) and assessed their baseline gene expression levels in purified blood cells from the BLUEPRINT consortium ^29^. Based on higher correlation with counts of the corresponding cell type and the higher expression in BLUEPRINT data, we concluded that eight out of the ten modules were strongly associated with specific blood cell types (Figure 2c and Figure S2), namely neutrophils, CD4+ T-cells, NK cells, CD8+ T-cells, monocytes, erythroblasts, macrophages and eosinophils.

The eQTLs modified by cell type proxies were enriched in cell-type-specific signaling pathways (Figure 2c, Table S8). For example, genes for which the *cis*-eQTL effects were particularly strong in erythroblasts (module 4) had erythrocyte-specific functions. They were also enriched in binding sites for transcription factors involved in erythrocyte development based on ENCODE ChIP-seq data (GATA1, TAL1, GATA2 and MafK, each with enrichment p-values ≤ 10^−5^) ^30–32^. For the other modules, we also identified specific transcription factors having established functions in the corresponding cell types (Table S9). Analysis of eQTL interactions with the blood cell counts measured confirmed the cell-type-dependent effects on neutrophils, lymphocytes, eosinophils and monocytes (Table S10). However, the comparison with BLUEPRINT expression data showed that our unbiased analysis also identified effects for cell types for which actual cell counts were not available (erythroblasts, CD4+ T-cells and NK cells/CD8+ T-cells), showing the usefulness of such an approach in datasets without comprehensive cell count measurements.

eQTLs that are lymphocyte-dependent and that show a stronger signal in B-, T- or NK cells are expected to replicate better within the in B-cell-derived LCL eQTL data generated by the Geuvadis consortium ^5^. We indeed found that replication of the lymphocyte-dependent eQTL effects is higher than our overall replication rate in Geuvadis (Table S11) (46% vs 38%, Fisher exact p-value = 0.005). In contrast, for our neutrophil-dependent eQTLs, the replication rate was similar to the overall replication rate (39% vs 38%, Fisher exact p-value = 0.58). Here, however, we observed that these neutrophil-dependent eQTLs are enriched for opposite allelic effects between our data and the Geuvadis results (27% opposite effects, compared to 14% for all overlapping eQTLs, Fisher exact p-value = 8.08 × 10^−5^). While the Geuvadis LCLs are not a perfect model for any of our lymphoid modules, our replication results do provide further evidence of the cell-type dependent nature of the identified context-dependent eQTLs.

The 10 proxy genes identified using the gene-level eQTLs also affected 4,216 exon-level eQTLs and 572 exon ratio QTLs. Gene function enrichment analysis on the exon-level and exon ratio QTLs showed results similar to that of eQTL genes (Table S8), indicating that the proxy genes do not solely represent the factors modulating gene-level eQTLs but also those that affect alternative splicing eQTLs.

### Risk factors for immune disorders show context-dependent effects

To demonstrate how eQTLs and their context-specific effects can help to better understand disease, we highlight *cis*-eQTLs for inflammatory bowel disease (IBD) and rheumatoid arthritis (RA). We performed clustering of the eQTL genes based on co-expression in our dataset.

These clusters consisted of genes expressed in the same cell types according to the BLUEPRINT data. Functional enrichment analyses of the eQTL genes revealed pathways relevant to the disease etiology. Moreover, the subset of context-specific eQTLs provides additional evidence for the cell types in which the disease risk alleles are most active. Examples are shown in Figures 3 (IBD), 4 (RA), S3 (other immune disorders), and Table S12.

**Figure 3.**
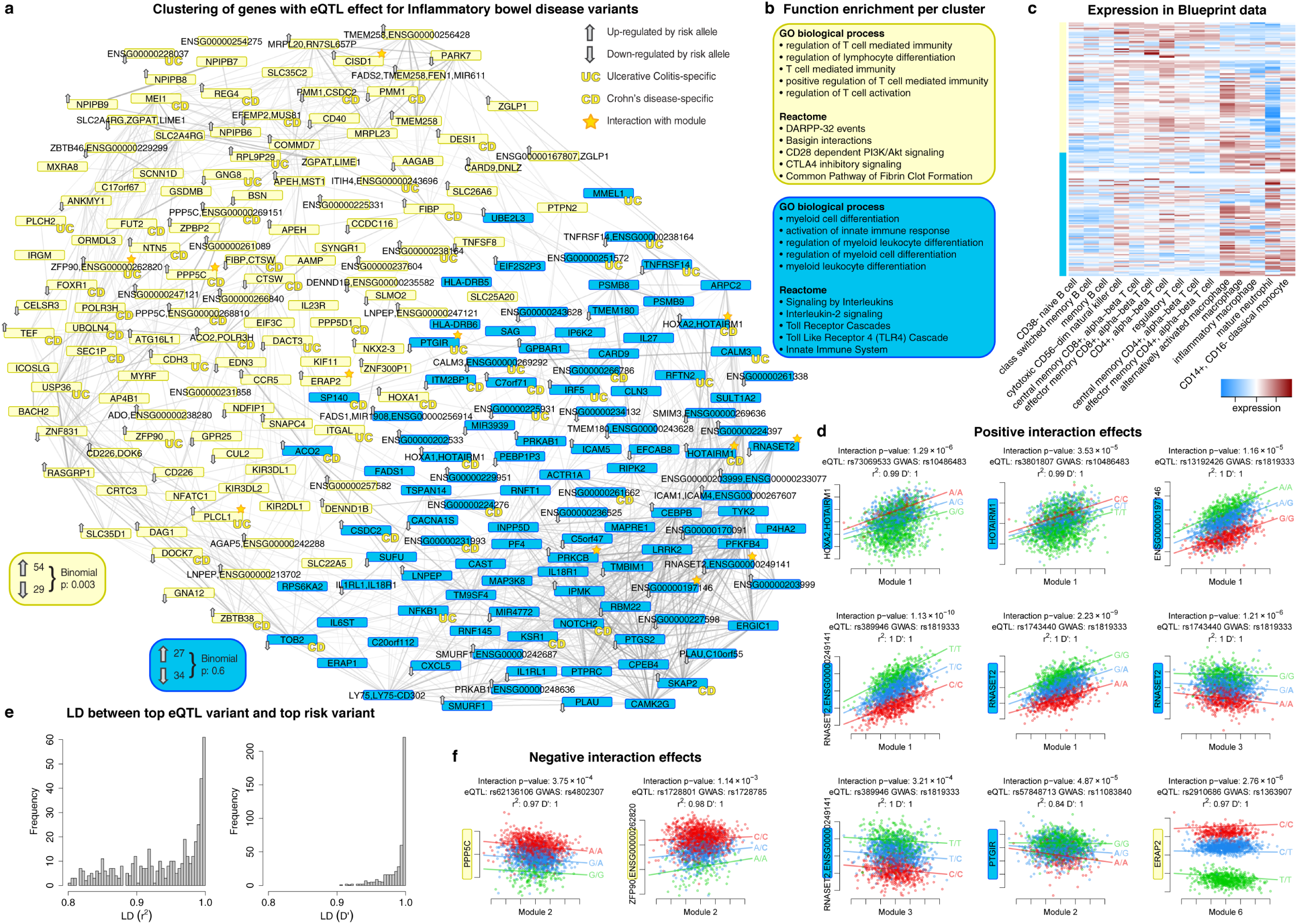
eQTLs associated with inflammatory bowel disease are predominantly active in neutrophils and T-cells. (A) The expression levels of 233 genes are regulated by 95 IBD loci. The gene-level eQTLs affected are indicated with arrows showing whether the risk allele increases or decreases expression. Genes without an arrow show exon QTL effects or polyA-ratio QTLs. Genes regulated by loci specific for ulcerative colitis (UC) or Crohn’s disease (CD) are marked accordingly (those without label are found in both). Genes regulated by an eQTL showing a significant interaction with one of our 10 modules are marked with a star. Clustering of the genes based on co-expression in our data revealed two clusters (colored yellow and blue). For the yellow cluster, we identified a significant overrepresentation of genes where the risk allele increases gene expression. (B) Gene function enrichment per cluster showed T-cell biology for the yellow cluster and neutrophil biology for the blue cluster. (C) Expression levels in the cell-sorted BLUEPRINT data show that the genes in the yellow cluster show higher expression in T-cells and the genes in the blue cluster show higher expression in neutrophils. (D) All positive eQTL interaction effects for IBD eQTLs. All positive interactions with module 1 (the neutrophil module) are by genes in the neutrophil cluster. (E) Distribution of LD (r^2^ and D’-values) between the top IBD variant and the top eQTL variant. (F) Two examples of negative interaction effects. Note: ENSG00000262820 has been removed in newer Ensembl versions. Figure 4. eQTLs associated with rheumatoid arthritis affect blood coagulation and NK- and B-cell biology. (A) We found 81 genes with eQTLs linked to RA. The gene level eQTL affected are marked with arrows indicating whether the risk allele increases or decreases expression. Genes without an arrow show exon QTL effects or polyA-ratio QTLs. Genes regulated by an eQTL showing a significant interaction with one of our 10 modules are marked with a star. Clustering of the genes based on co-expression revealed three clusters (colored yellow, aquamarine and blue). (B) Positive interaction for eQTL effects of RA loci. Genes in the blue NK-cell cluster show interactions for module 3 (the NK/CD8+ T-cell module) and genes in the aquamarine B-cell cluster show interaction in module 9 (the B-cell module). (C) Gene function enrichment per cluster showed B-cell biology for the aquamarine cluster and NK-cell pathways for the blue cluster. The yellow cluster shows different types of pathways not specific to a cell type. (D) Expression levels in the cell-sorted BLUEPRINT data confirm the B-cell andNK-cell annotations of the aquamarine and blue clusters. (E) Distribution of LD (r^2^ and D’-values) between the top IBD variant and the top eQTL variant.

For IBD we included the loci reported by a recent multi-ethnic meta-analysis ^33^. Of the 232 top SNPs reported in this meta-analysis, 95 loci (41%) are in strong LD (r^2^ ≥ 0.8) with a top eQTL SNP (median r^2^ = 0.96 and median D’ = 0.996). Importantly, 42 risk variants are in perfect LD with the top eQTL SNP (Figure 3e), which was proportionally much higher compared to non-immune traits (Figure 1c). In total, eQTL SNPs that are proxies for these 95 IBD loci affected expression of 233 genes (Figure 3a).

Clustering of these 233 genes revealed a set of 123 genes mainly related to T-cell biology and a set of 110 genes specific to neutrophils and toll-like receptor signaling (Figure 3b and 3c, see Table S13 for full enrichment analysis results), confirming existing knowledge of the cell types relevant in IBD ^20,21^. There was a significant imbalance in the direction of regulation within the T-cell cluster: 54 genes were up-regulated by the IBD risk allele whereas only 29 were down-regulated (binominal test p-value: 0.003), suggesting increased T-cell activity in IBD.

Incorporation of exon level eQTLs can be used to better understand gene level eQTLs. For instance, rs727088 lowered *CD226* expression. Zooming into the exon QTL effects of this gene, we found two exons that were strongly up-regulated, both of them unique to a non-sense mediate decay (NMD) transcript (as annotated by Ensembl). The other exons showed down-regulation by the risk allele, suggesting that a shift to the NMD isoform is lowering overall gene expression levels (Figure S4).

Twelve IBD-linked genes demonstrated context-dependent eQTL effects (Table S12). Five of these eQTLs were strongest in neutrophils (positive interaction score for module 1) and the genes containing these eQTLs were present in the neutrophil cluster (Figure 3d). We also observed negative interactions, where the effect becomes smaller in a specific module, *e.g*. the eQTL effect of rs1728801 regulating *ZPF90* (Figure 3f), a gene that is known to be important in T-helper cells ^36^. We found that carriers of the protective allele show increased expression of *ZPF90* in CD4+ T-cells, while the risk allele carriers show consistently high expression levels independent of CD4+ T-cells. This indicates that while *ZFP90* expression normally is restricted to T-cells this does not apply to carriers of the risk allele which have high expression levels in multiple cell-types, potentially activating T-cell specific pathways in other cells.

For RA, 81 eQTL genes were found that cluster into three groups (Figure 4a). The median r^2^ between the top eQTL SNPs and the top GWAS hits is 0.95, the median D’ is 0.996 and 24 variants are in perfect LD. The genes in two clusters showed high expression levels in B-cells and NK/T-cells, respectively (Figure 4d). This was supported by the gene function enrichments (Figure 4c). The genes in the third cluster were highly expressed in macrophages, neutrophils and monocytes and were related to platelet activation and coagulation. RA patients are known to be at increased risk for venous thromboembolism ^25^ and the importance of blood coagulation factors in RA pathogenesis has recently been established ^26^. The eQTL results link risk loci and genes to this symptom.

**Figure 4.**
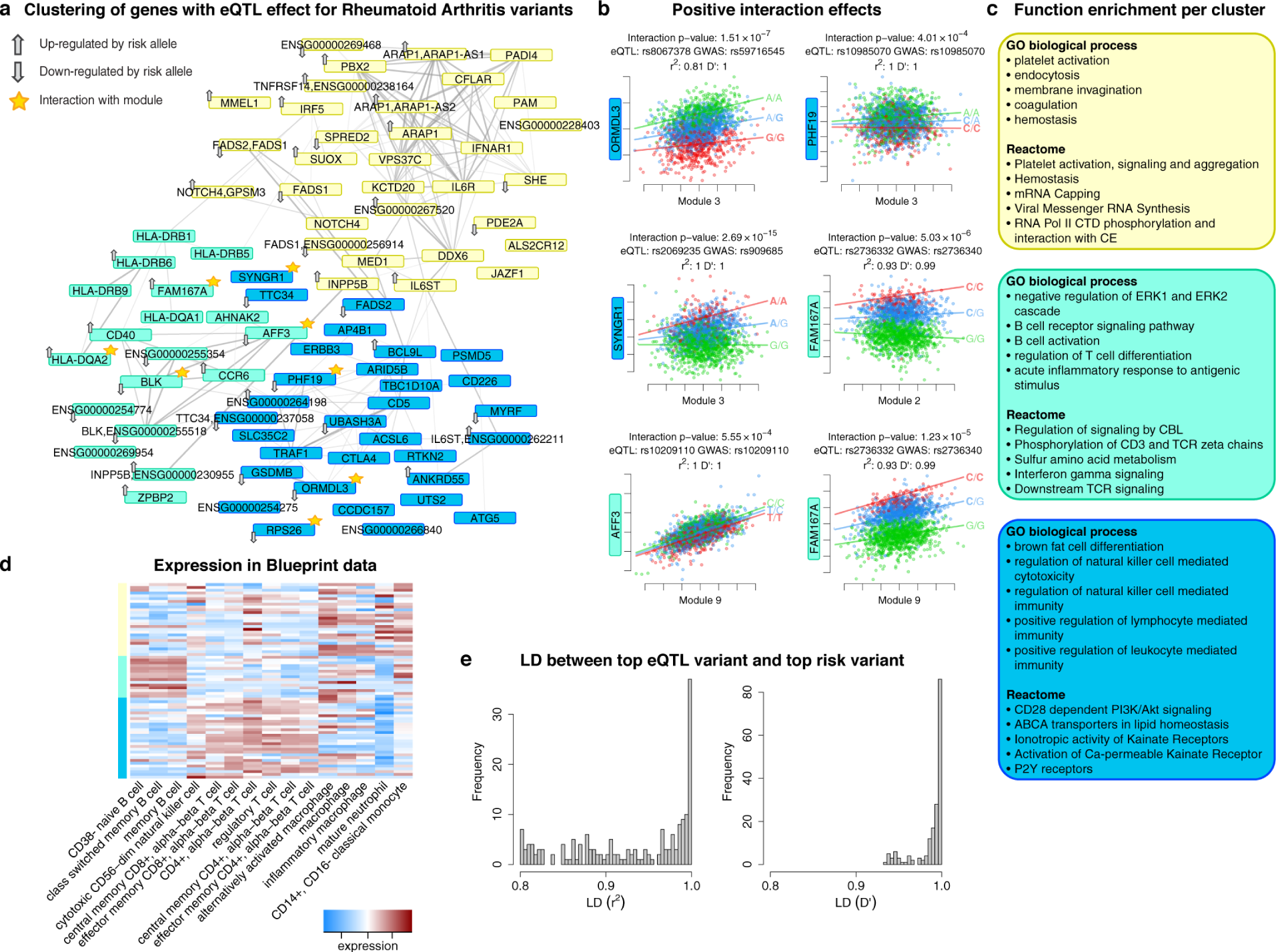
eQTLs associated with rheumatoid arthritis affect blood coagulation and NK- and B-cell biology. (A) We found 81 genes with eQTLs linked to RA. The gene level eQTL affected are marked with arrows indicating whether the risk allele increases or decreases expression. Genes without an arrow show exon QTL effects or polyA-ratio QTLs. Genes regulated by an eQTL showing a significant interaction with one of our 10 modules are marked with a star. Clustering of the genes based on co-expression revealed three clusters (colored yellow, aquamarine and blue). (B) Positive interaction for eQTL effects of RA loci. Genes in the blue NK-cell cluster show interactions for module 3 (the NK/CD8+ T-cell module) and genes in the aquamarine B-cell cluster show interaction in module 9 (the B-cell module). (C) Gene function enrichment per cluster showed B-cell biology for the aquamarine cluster and NK-cell pathways for the blue cluster. The yellow cluster shows different types of pathways not specific to a cell type. (D) Expression levels in the cell-sorted BLUEPRINT data confirm the B-cell and NK-cell annotations of the aquamarine and blue clusters. (E) Distribution of LD (r2 and D’-values) between the top IBD variant and the top eQTL variant.

Eight RA-associated eQTLs were context dependent (Table S12). In concordance with their gene expression patterns, six of these demonstrated stronger eQTL effects in T- or B-cells (Figure 4b). *FAM167A* demonstrated an interaction with both module 9 (B-cells) and module 2 (CD4+ T-cells). Interestingly, we observe that for carriers of the protective allele there is no correlation between CD4+ T-cells and the expression levels of *FAM167A*, whereas for carriers of the risk allele we do see up-regulation of this gene in CD4+ T-cells.

We obtained similar results for celiac disease (Figure S3a), type 1 diabetes (Figure S3b), multiple sclerosis (Figure S3c) and systemic lupus erythematosus (Figure S3d).

### Modules not associated with cell types

We found that most modules represented different cell types, except module 6 and 7. Although module 6 could not be attributed to a clear biological process, module 7 contains many genes that are known to be involved in the type I interferon response. Pathway enrichment of the 51 module 7 proxy genes, which were positively correlated with the expression of the top proxy gene *SP140*, revealed that many of these genes are involved in the type I interferon response to viral infection. The other 48 genes that correlated negatively with *SP140* are involved in anti-bacterial response and inflammation (Figure 5a). The positively correlated genes are enriched for up-regulated genes upon rhinovirus stimulation ^16^ (Fisher exact p-value 1.14 × 10^−9^), in line with their involvement in the type I interferon response. In contrast, the negatively correlated genes are enriched for genes up-regulated upon LPS stimulation (Fisher exact p-value 0.02) and interferon-gamma stimulation (Fisher exact p-value 8.72 × 10^−4^) ^15^, supporting the anti-bacterial function of these genes (Table S14). Likewise, the affected eQTL genes can be divided into two groups: those that were positively correlated with *SP140* expression and those with negative correlations to *SP140*. Similarly to the proxy genes, the most significantly enriched pathway for the positively correlated eQTL genes was ‘response to type I interferon’ (Figure 5b). Gene annotations from the interferome database ^39^ confirmed that the up-regulated eQTL genes are indicative of a type I, but not of a type II interferon response (Figure 5c). Type I interferon signaling is activated in a viral response and type II interferon signaling is activated upon bacterial response ^40^. Both type I and type II interferon signaling result in binding of heterodimers of the STAT1 transcription factor. Unique to type I interferon is that STAT1 forms a complex with STAT2 and IRF9, resulting in the activation of viral response genes. STAT3 activation is also unique to the type I response, resulting in the down-regulation of inflammatory pathways ^41^. The eQTLs were enriched for STAT1 (p-value = 4.82 × 10^−04^), STAT2 (3.12 × 10^−04^) and STAT3 (4.72 × 10^−05^) binding sites (based on ENCODE ChIP-seq experiments) (Table S9). Motif enrichment analysis ^42^ on the 25 bp flanking regions of the eQTL SNPs confirmed the enrichment of STAT-binding motifs (Wilcoxon rank-sum test, p-value = 9.61 × 10^−05^). In support of the modifying effects of viral cues on this set of eQTLs, eQTL genes that have recently been reported as rhinovirus-response QTLs ^16^ typically have higher interaction z-scores for module 7 than other eQTL genes (Wilcoxon p-value = 0.02). We therefore conclude that the effect of these 145 eQTL genes is dependent on stimulation with type I interferon.

**Figure 5.**
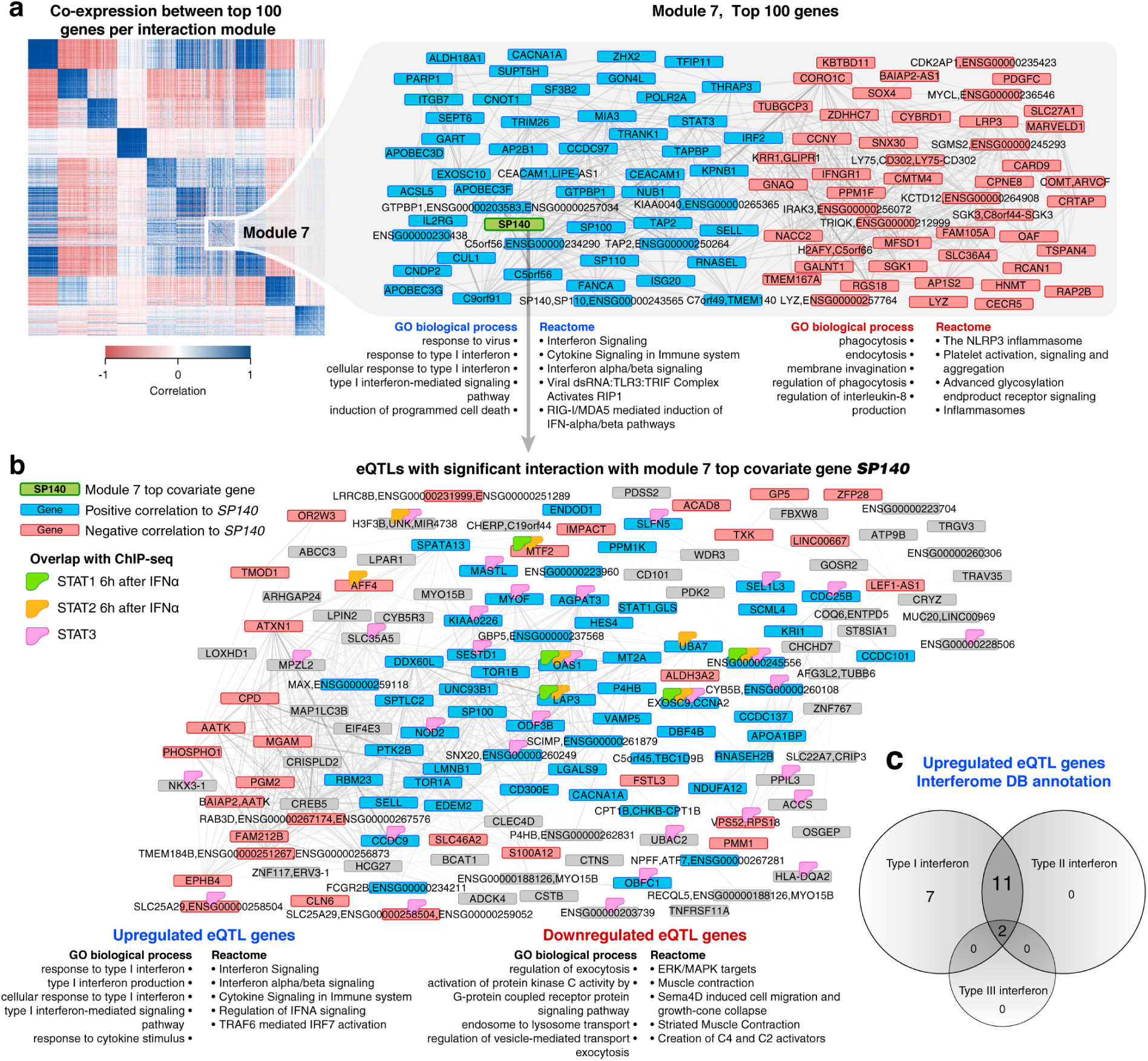
eQTLs modified by type I interferon signaling. (A) The top 100 covariate genes for module 7 are clustered based on expression levels, revealing two distinct clusters of genes. Genes positively correlated with the top covariate *(SP140)* are indicated in blue and those negatively correlated with *SP140* in red. Enrichment analysis of these two clusters show distinct biology: the up-regulated genes are enriched for type I interferon response and response to viruses whereas the down-regulated genes indicate an anti-bacterial inflammatory response. (B) The eQTLs affected by module 7 can also be divided into those genes positively or negatively correlated with *SP140* expression. The significantly positively correlated eQTL genes are also enriched for type I interferon response, whereas the negatively correlated eQTL genes do not show strong enrichment for biological functions. (C) Interferome DB annotation of the up-regulated eQTL genes confirmed their role in type I (and not type II or III) interferon signaling.

### Regulatory network discovery

Each of the aforementioned ten modules demonstrated effects on many (>120) eQTLs. However, some other factors may also exist that affect more limited numbers of eQTLs. To identify these, we first corrected the expression data for the 10 module interaction effects and then ascertained for each gene-level eQTL whether the eQTL effect size was significantly dependent on the expression of any other gene. This resulted in the identification of an additional set of 901 context-dependent eQTL genes (FDR ≤ 0.05). Of these eQTL interactions, 113 could be replicated in Geuvadis LCLs (FDR ≤ 0.05, 94% with the same interaction direction) (Table S15). These LCLs are derived from a single cell type so any interaction effect that replicates is unlikely due to cell type specific eQTL effects, but likely reflect an external stimulation or activation of core biological processes. A few of these context-dependent eQTLs enable inference of regulatory networks:

An example is a *cis*-eQTL effect on the lipid biosynthesis gene *FADS2* that is modified by the expression of the sterol regulatory element binding transcription factor *SREBF2* (p-value = 4.1 × 10^−14^, p-value in Geuvadis = 0.002) (Figure 6a and 6b). The eQTL SNP was in close proximity to an SREBF2 binding site (ENCODE ChIP-seq data, Figure 6c) and it is therefore likely that the SNP modifies the affinity of the *FADS2* promoter for SREBPF2. *SREBF2* showed a significant negative correlation to HDL cholesterol (Pearson r = -0.18, p-value = 5.1 × 10^−6^) and a positive correlation to lymphocyte percentage (Pearson r = 0.19, p-value = 1.6 × 10^−6^). A partial correlation analyses revealed that the correlation to HDL cholesterol is independent of the correlation to the lymphocyte percentage (Pearson r on residuals of HDL after correcting for lymphocytes: -0.17, p-value = 2.7 × 10^−5^), showing that the correlation to HDL is not driven by cell type composition. We propose a model where extracellular (HDL) cholesterol levels modify SREBF2 binding to the *FADS2* promoter, which, in turn has effects on the expression of FADS2 and the lipid unsaturase activity in the cell.

**Figure 6.**
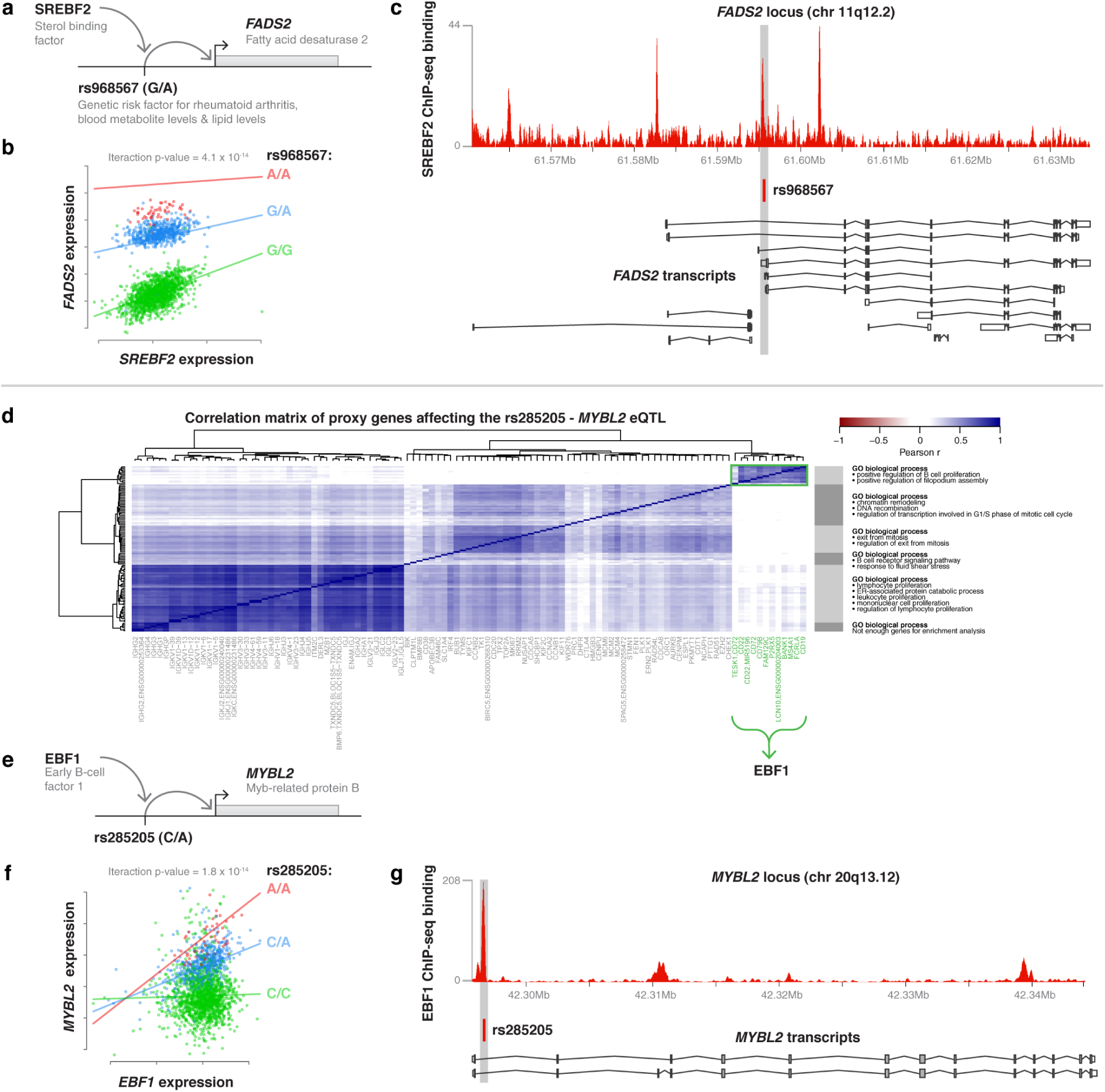
Context-dependent eQTL effects not present in the 10 modules. (A & B) *SREBF2* expression modulates the eQTL effect on *FADS2*. (C) The eQTL SNP affecting *FADS2* expression is located in an ENCODE ChIP-seq peak of *SREBF2* binding. (D) Heatmap of the co-expression of 109 proxy genes that affect the eQTL effect on *MYBL2* expression. The different clusters show pathway function enrichments related to proliferation and cell-cycle. (E) Regulation of *MYBL2* by the different cell-cycle clusters is likely modulated via EBF1 and rs285205. (F) Interaction plot when using *EBF1* as proxy gene (G) EBF1 binds directly over the *MYBL2* eQTL SNP rs285205.

We found 109 genes that alter the rs285205 *cis*-eQTL effect on *MYBL2*, which encodes a known transcription factor that controls cell division and has a known tumor suppressing function ^43^. Co-expression clustering based on these 109 proxy genes revealed six clusters (Figure 6d). Gene function enrichment analyses on the genes in these clusters revealed that all were related to proliferation or cell cycle checkpoints. Interestingly, only one cluster increased the magnitude of the *MYBL2* eQTL effect in contrast to the other clusters, which all repressed this eQTL. This eQTL activating cluster was strongly enriched for "positive regulation of B cell proliferation” (p-value = 1 × 10^−7^), and the strongest proxy gene in this cluster was *FCRLA*, which is known to be highly expressed in proliferating B-cells residing in the germinal center of the lymph nodes (centroblasts) ^44^. *FCRLA* is also reported to be highly expressed in lymphoblastoid cell lines, which are in a highly proliferative state ^45^. However, in our analysis we had initially only considered genes that where expressed in each of our individuals (see methods), and therefore had not studied low-abundant transcription factor genes. When also including these genes, we observed this cluster of genes is strongly co-expressed with EBF1, a transcription factor that drives B-cell differentiation and proliferation, suggesting that EBF1 might drive the eQTL interaction effect for *MYBL2*. This was indeed confirmed using ENCODE ChIP-seq data in LCLs: we observed strong binding of EBF1 at rs285205 (Figure 6g). EBF1 is a known player in B-cell differentiation and proliferation and positively correlated to both *MYBL2* (r = 0.11, p-value = 6.99 × 10^−7^) and *FCRLA* (r = 0.8, p-value ≤ 2.2 × 10^−16^). Finally, testing *EBF1* as a proxy gene directly also revealed a significant interaction (p-value = 1.8 × 10^−14^) effect (Figure 6f). As such EBF1 influences *MYBL2* gene expression, but because of its binding at SNP rs285205, this SNP likely affects the binding affinity of EBF1. In the Geuvadis dataset these *MYBL2* interactions are not significant, however 74% were in the same direction (binominal p-value = 2.2 × 10^−5^).

## Discussion

Using whole blood RNA-seq data we greatly expanded the catalog of SNPs that have a known regulatory function. Our study on 2,116 individuals allowed identification of many independent genetic effects influencing expression levels. We observe co-localization of one third of the disease- and trait-associated variants with eQTL signals, and we observed significantly more overlap for traits related to blood (immunological disorders/metabolic levels) as compared to other traits such as height and BMI, and thus arrive at different conclusions as compared to a recent paper that observed that blood eQTL and IBD variants do not significantly more often co-localize as compared to chance ^46^. However, since that analysis used earlier array-based blood eQTL results that had not been imputed to recent reference panels, this suggests that earlier eQTL studies either have lacked sufficient statistical power to identify eQTLs (i.e. limited sample size and/or the use of microarrays instead of RNA-seq) or were unable to fine-map these associations (i.e. imputation to older reference panels). Additionally, the presence of multiple independent regulatory variants might have been missed; while the regulatory effects of nearly 500 GWAS hits were not in strong LD with our top eQTL SNPs, they were in strong LD with our secondary, tertiary or even higher order eQTLs (Figure 1a, Figure S1). Moreover, approximately 10% of the GWAS hits did not affect the regulation of overall gene expression levels, but had an impact on mRNA processing because they were found only in the exon, exon ratio or polyA-ratio QTL analyses (Figure 1b, Table S6, examples in Supplementary Material online).

To gain a better understanding of the biology behind these regulatory variants, we identified 2,743 context-dependent eQTLs (1,842 in the first 10 modules and 901 in the remainder) and identified many of the determinants that modify these eQTLs. These provide further insight into the cell types in which the genetic risk factors are regulating gene expression and the regulatory networks in which they participate, further refining our findings on GWAS risk loci. Unlike other approaches (^15, 16, 20^), our method does not rely on any prior knowledge or assumptions on differences in cell type composition or naturally occurring stimulations acting on our whole tissue data.

We also observed various examples where eQTLs are influenced by transcription factors that bind to the eQTL SNP. This includes rs968567 that strongly influences *FADS2* gene expression and depends on SREBF2 activity, but which also increases risk for rheumatoid arthritis, blood metabolite levels and lipid levels. With future increases in sample-size we expect it might also become possible to observe gene x environment interaction effects for this SNP on these (disease) phenotypes as well.

Our method can easily be applied to other tissues and we expect that many more cell-type-dependent eQTLs will become detectable in such datasets once sample-sizes become sufficiently large. This will likely lead to the identification of even more unanticipated intrinsic factors and external stimuli that modify the downstream effects of genetic risk factors. As such our approach complements perturbation experiments to gain better insight in regulatory networks and their stimuli.

## Acknowledgements

This work was performed within the framework of the Biobank-Based Integrative Omics Studies (BIOS) Consortium funded by BBMRI-NL, a research infrastructure financed by the Dutch government (NWO 184.021.007). Samples were contributed by LifeLines (http://lifelines.nl/lifelines-research/general), the Leiden Longevity Study (http://www.healthy-ageing.nl; http://www.leidenlangleven.nl), the Rotterdam studies (http://www.erasmus-epidemiology.nl/rotterdamstudy) and the CODAM study (http://www.carimmaastricht.nl). We thank the participants of all aforementioned biobanks and acknowledge the contributions of the investigators to this study (Supplemental Acknowledgements). This work was carried out on the Dutch national e-infrastructure with the support of SURF Cooperative and the Groningen Center for Information Technology (Strikwerda, W. Albers, R. Teeninga, H. Gankema and H. Wind) and Target storage (E. Valentyn and R. Williams). Target is supported by Samenwerkingsverband Noord Nederland, the European Fund for Regional Development, the Dutch Ministry of Economic Affairs, Pieken in de Delta and the provinces of Groningen and Drenthe.

## Data availability

All results can be queried using our dedicated QTL browser:

http://genenetwork.nl/biosqtlbrowser/. Raw data was submitted to the European Genome-phenome Archive (EGA, accession number EGAS00001001077).

## Author contributions

BTH, PACtH, JBJvM, AI, RJ and LF formed the management team of the BIOS consortium. DIB, RP, JvD, JJH, MMJVG, CDAS, CJHvdK, CGS, CW, LF, AZ, EFG, PES, MB, JD, DvH, JHV, LHvdB, CMvD, AH, AI, AGU managed and organized the biobanks. JBJvM, PMJ, MV, JvR and NL generated RNA-seq data. HM, MvI, MvG, WA, JB, DVZ, RJ, PvtH, PD, MV, IN, MaS, PACtH, BTH and MM were responsible for data management and the computational infrastructure. DVZ, PD, MV, MvI, FvD, MvG, WA, MJB, HJW, SMK, JL, MAS, PACtH and LF performed the data analysis. DVZ, PD, PACtH and LF drafted the manuscript.

